# An Antibody to RGMa Promotes Regeneration of Cochlear Synapses after Noise Exposure

**DOI:** 10.1101/2020.07.01.183269

**Authors:** Jerome Nevoux, Mihaela Alexandru, Thomas Bellocq, Lei Tanaka, Yushi Hayashi, Takahisa Watabe, Hanae Lahlou, Kohsuke Tani, Albert S.B. Edge

## Abstract

Auditory neuropathy is caused by the loss of afferent input to the brainstem via the components of the neural pathway comprising inner hair cells and the first order neurons of the spiral ganglion. Recent work has identified the synapse between cochlear primary afferent neurons and sensory hair cells as a particularly vulnerable component of this pathway. Loss of these synapses due to noise exposure or aging results in the pathology identified as hidden hearing loss, an initial stage of cochlear dysfunction that goes undetected in standard hearing tests. We show here that repulsive axonal guidance molecule a (RGMa) acts to prevent regrowth and synaptogenesis of peripheral auditory nerve fibers with inner hair cells. Treatment of noise-exposed animals with an anti-RGMa blocking antibody regenerated inner hair cell synapses and resulted in recovery of wave-I amplitude of the auditory brainstem response, indicating effective reversal of synaptopathy.

## INTRODUCTION

Repulsive guidance molecule (RGM) has an important role in many fundamental process, including cell migration, differentiation and apoptosis both during and after development of different organs (Siebold et al., 2017). RGM was discovered through studies on developing axons during embryogenesis (Monnier et al., 2002); since its discovery, four different forms of this soluble or membrane bound molecule have been reported in vertebrates: RGMa, RGMb (or DRAGON), RGMc (or hemojuvelin) and RGMd (only in fish) (Demicheva et al., 2015; Siebold et al., 2017).

RGMs are membrane associated glycosylphosphatidylinositol (GPI)-linked proteins that contain an N-terminal signal peptide, an RGD motif, and a partial von Willebrand type D structural domain (Severyn et al., 2009). RGMs bind the type 1 transmembrane protein neogenin1, which controls axon guidance and neuronal survival effects. The binding of two RGM molecules to the juxtamembrane fibronectin type III (FNIII) domains of two neogenin1 receptors brings them into close association. The resulting intracellular dimerization acts as a mechanism of signal transduction through the plasma membrane. In addition, the RGMs are co-receptors for bone morphogenetic proteins (Siebold et al., 2017).

RGM has key functions in the nervous system. The role in the optical system is well known but a lot is still to discover in the auditory system. Blocking the interaction between RGM and neogenin1 with anti-RGMa antibodies prevents proapoptotic and neurite growth inhibitory effects. Using these antibodies in brain or axonal injury leads to neuronal and synaptic regeneration with functional recovery (Demicheva et al., 2015).

In previous studies we found that an antibody against RGMa increased the innervation of postnatal organ of Corti by spiral ganglion neurons (Brugeaud et al., 2014). These experiments were performed as an *in vitro* model for the reinnervation of auditory neurons by embryonic stem cells in an attempt to devise stem cell based treatments for auditory neuropathy (Corrales et al., 2006; Martinez-Monedero et al., 2006; Tong et al., 2013), the loss of auditory neurons that occurs in genetic diseases as well as age and noise related insults to the cochlea.

The cochlear sensory synapses are glutamatergic synapse with glutamate as the neurotransmitter of inner hair cells. Kainic acid (KA), an analog of glutamate, has a depolarizing effect on neurons but can also be neurotoxic (Coyle, 1983). Pujol et al. reported the severe swelling of auditory dendrites in the organ of Corti after a local injection of 1 nmol KA (Pujol et al., 1985). This result is consistent with previous ultrastructural analyses of KA neurotoxicity in other neuronal tissues. This type of damage is another form of auditory neuropathy in which the cell bodies of the neurons remain but the synapses are disrupted, and recent evidence suggests that this type of damage occurs as the primary lesion in sensorineural hearing loss (Kujawa and Liberman, 2006, 2009; Sergeyenko et al., 2013; Wu et al., 2019). Here we tested whether the inhibition of RGMa would aid in the regeneration of these synapses both in an *in vitro* model of kainate excitotoxicity and i*n vivo* after noise-induced damage to the synapse. The results suggest that indeed the inhibition of RGMa is an effective treatment for the synaptic loss and may therefore be a viable approach to the treatment of patients with this pathology

## METHODS

### Organ of Corti culture

The cochlea from postnatal day (P)4 to P6 mice of both sexes were dissected according to the protocol described (Parker et al., 2010). The membranous labyrinth was separated from the bone, and the stria vascularis and spiral ligament were carefully removed. Reissner’s membrane and tectorial membrane were removed with fine forceps with the organ of Corti and spiral ganglion maintained intact. The middle region of the cochlea was cultured in a well on a cover glass coated with 1:1 poly-L-ornithine (0.01%; Sigma-Millipore #P4957, USA) and laminin (50 μg/mL; BD Biosciences, USA). The explants were cultured in N2 and B27-supplemented DMEM, 1% HEPES solution (#H0887, Millipore Sigma, USA), 1:1000 Ampicillin, 1:300 Fungizone at 37°, 6.5% CO2 for up to 6 days.

### Noise exposure and kainic acid model

Male mice CBA/CaJ were received from Jackson laboratories at 7 weeks old and exposed at the age of 8 weeks, awake and unrestrained, to octave-band noise (8–16 kHz) for 2 h at 98 dB sound pressure level (SPL) in a reverberant sound-exposure box. Mice were placed within acoustically transparent wire cages on a rotating platform. The noise waveform was generated digitally using an h-order Butterworth filter, amplified through a power amplifier (Crown D75A) and delivered by a loudspeaker (JBL2446H) coupled to an exponential horn in the roof of the box. Sound levels were verified in the center of the cage with a 1/4′′ Bruel and Kjaer condenser microphone before each exposure and varied by less than 1 dB in the cage space during the whole exposure.

To reproduce the excitotoxic trauma *in vitro*, we adapted a rat model of excitatory cochlear synaptopathy (Wang and Green, 2011) to the mouse explants. We exposed organ of Corti explants, after 24 h in culture to a solution of 0.4 mM kainic acid (#ab120100, Abcam, USA) diluted in culture medium for 2 h.

### Cochlear testing

ABRs and DPOAEs were recorded just before and 24 h after noise trauma, then just before sacrifice 3 weeks after noise exposure. The animals were anesthetized with an intraperitoneal injection of ketamine (100 mg/kg) and xylazine (20 mg/kg) and placed in an acoustically and electrically shielded room maintained at 32 °C. Acoustic stimuli were delivered through a custom acoustic system consisting of two miniature dynamic earphones used as sound sources (CDMG15008-03A, CUI) and an electret condenser microphone (FG-23329-PO7, Knowles) coupled to a probe tube to measure sound pressure near the eardrum. Custom LabVIEW software controlling National Instruments 24-bit soundcards (6052E) generated all ABR/ DPOAE stimuli and recorded all responses.

For ABRs, stimuli were 5 ms tone pips (0.5 ms cos2 rise-fall) at frequencies from 5.66 to 45.25 kHz (in half-octave steps) delivered in alternating polarity at 35/s. Electrical responses were collected via needle electrodes at the vertex, the pinna and a ground reference near the tail, amplified 10,000X with a 0.3–3 kHz passband, and averaged with 512 responses at each SPL. Responses were collected for stimulus levels in 5-dB steps from 10 dB below threshold up to 90 dB SPL. ABR threshold was defined as the lowest sound level at which a reproducible waveform could be observed. When there was no detectable response at 90 dB SPL, threshold was defined as 95 dB. Wave-I amplitude was defined as the difference between the average of the 1-ms pre-stimulus baseline and the wave-I peak (P1), after additional filtering to remove low-frequency baseline shifts.

For DPOAEs, the cubic distortion product 2f1 − f2 was measured in response to primaries f1 and f2 (frequency ratio f2/f1 = 1.2, and level ratio L1 = L2 + 10), where f2 varied from 5.66 to 45.25 kHz in half-octave steps. Primaries were swept in 5 dB steps from 10 to 80 dB SPL (for f2). The DPOAE at 2f1 − f2 was extracted from the ear canal sound pressure after both waveform and spectral averaging. DPOAE threshold was computed by interpolation as the primary level (f1) required to produce a DPOAE of 0 dB SPL.

### Surgical techniques and RGMa treatment

The surgery was performed 1 week after noise trauma at the age of 9 weeks old. CBA/CaJ male mice (25-35 g, 9 weeks old) were divided into 3 groups: group A (no noise trauma, no treatment, negative control group), group B (noise trauma and rabbit IgG anti-rat RGMa (#28045, IBL, Japan)), group C (noise trauma and rabbit IgG, control for surgery (#SC2027, Santa Cruz, USA). The mice were anesthetized with intraperitoneal (IP) injection of ketamine (100 mg/kg) and xylazine (20 mg/kg). The right bulla was exposed via a dorsal lateral approach to administer 10 μL of IgG anti-RGMa solution (10 μg/mL diluted in PBS) or IgG solution (10 μg/mL diluted in PBS) onto the intact right round window membrane (RWM), using a microliter syringe (NanoFil-100 μL, WPI, USA) mounted with a 36G microneedle (#NF33-36BV, WPI, USA) under a surgical microscope. The left ear was used as control. After local administration, the RWM was kept in a horizontal position for 1 hour. The wound was closed with resorbable suture material. All experimental procedures were approved by the Institutional Animal Care and Use Committee of the Massachusetts Eye and Ear Infirmary and conducted in accordance with the NIH Guide for the Care and Use of Laboratory Animals.

In the *in vitro* model, after KA treatment, the explants were maintained for up to 5 days in 80 μL of control culture medium, culture medium containing anti-RGMa antibody at 10 μg/mL (#28045, IBL, Japan), or IgG at 10 μg/mL (normal rabbit IgG, #SC2027 Santa Cruz, USA), changed daily. The cultures were then fixed and prepared for immunohistochemistry.

### Cochlear immunohistochemistry

For the whole-mount (Shi et al., 2013), the mice were anesthetized with intraperitoneal (IP) injection of ketamine (100 mg/kg) and xylazine (20 mg/kg) and transcardially perfused with 4% paraformaldehyde (PFA). The cochlea was harvested and post-fixed for 2 h in PFA 4% at room temperature. It was then decalcified in 0.2 M EDTA pH 7.4 for 2 days at room temperature. The cochlea was dissected into 6 pieces and incubated in 0.3% Triton X-100 and 5% normal horse serum at room temperature for 1 h for permeabilization and blocking.

Cochlear pieces were mounted using fluorescent mounting medium (Dako, Denmark) and coverslips. Organ of Corti cultures were fixed with 4% paraformaldehyde for 15 min and blocked with the same solution. Eight-week old CBA/Caj strain mouse cochlea was used for neogenin1 and RGMa immunostaining. The cochlea was processed and embedded in OCT compound, (Tissue-Tek Sakura, Finetek, USA) and cut using a cryomicrotome (#818, Leica, Germany). Sections were frozen and subjected to antigen-retrieval using citrate buffer (#ab93678, Abcam,USA) before blocking and staining.

Immunostaining was done using the following primary antibodies: a rabbit anti-RGMa polyclonal antibody (#28045, IBL, USA), a mouse anti-neogenin purified monoclonal antibody (#AF1079, R&D, USA), a mouse (IgG1) anti-CtBP2 at 1:200 (#612044, BD Transduction Labs), a mouse (IgG2a) anti-GluA2 (Glutamate receptor subunit A2) at 1:2000 (#MAB397, Millipore) a chicken anti-neurofilament (AB5539, Millipore, USA) and a rabbit anti-myosin7a (#25-6790, Proteus Biosciences, USA).

Secondary antibodies were all used at 1:500 and were: Alexa Fluor 450-conjugated goat anti-rabbit (#A31556, Life Technologies, USA); Alexa Fluor 488-conjugated goat Mouse (IgG2) (#A21131, Life Technologies, USA); Alexa Fluor 568-conjugated goat anti-Mouse (IgG1) (#A21124, Life Technologies, USA); Alexa Fluor 647-conjugated goat anti-chicken (#A21449, Life Technologies USA).

### In situ hybridization

For RNA in situ hybridization, we used RNAscope Multiplex Fluorescent Reagent Kit (Advanced Cell Diagnostics). We prepared cryosections from P4 mouse cochlea and post-fixed the slides with 4% paraformaldehyde at room temperature for 40 min. Slides were washed twice with phosphate-buffered saline at room temperature for 5 min and soaked in 100% ethanol and hybridized according to the manufacturer’s instructions. After these procedures, the samples were stained for GFP, f-actin, and DAPI.

### Synapse counts

For cochlear whole mounts, confocal z-stacks were obtained from the inner hair cell area using a high-resolution glycerin-immersion objective (63×, 1.3 N.A.) and 3.10 × digital zoom with a 0.25 μm z-spacing on a Leica SP8 confocal microscope. For each stack, the z-planes images included all synaptic elements in the x-y field of view, which encompassed ∼7-8 inner hair cells. To identify and measure pre-synaptic ribbons and postsynaptic glutamate receptor patches, confocal z-stacks were exported to image-processing software (Amira, Visage Imaging, Thermo Fisher Scientific, USA). Synaptic puncta were identified using the connected components feature, which records the volume and x,y,z position of each synaptic element, defined as discrete volumes of at least 10 contiguous voxels in which intensity values in the selected channel are greater than an arbitrary criterion (usually set at 60 on a 0– 255 8-bit scale). For each identified punctum, custom software accessed the z-stack, extracted a voxel cube (1 μm on a side) centered on its x,y,z, coordinate, computed the maximum projection of this cube, and displayed it as a thumbnail image. Ribbon positions were confirmed manually so their total number was counted, and then divided by the number of inner hair cells, giving a mean number of ribbon synapses per cell.

### Statistical analysis

Statistical analyses were conducted using Prism (GraphPad Software, San Diego, CA, USA) for analysis of variance (ANOVA). Threshold shift data are presented as means ± standard errors of the mean (SEM).

## RESULTS

### Localization of neogenin1 and RGMa in the adult mouse cochlea

Neogenin1 is a guidance receptor expressed in the growth cone of neurons, and its binding to RGMa induces growth cone collapse, axon repulsion and neurite growth inhibition (Siebold et al., 2017). Their interaction is an important part of the regulation of the retinotectal path finding *in vivo* in the chick, allowed by different gradients of neogenin1 and RGMa.

RGMa and its receptor neogenin1 were shown to be expressed in the embryonic, newborn, and adult mouse cochlea (Brugeaud et al., 2014). Neogenin1 was localized to spiral ganglion neurons at P4, whereas RGMa could only be detected at the RNA level, by RT-PCR and *in situ* hybridization (Brugeaud et al., 2014).

Here we detected RGMa by immunostaining of the mouse cochlea (Fig. 1A), which showed RGMa protein juxtaposed to the afferent nerve fibers as they projected to inner hair cells (Fig. 1A). RGMa surrounded the afferent nerve endings at the inner hair cells. Since RGMa is a GPI-linked protein that becomes soluble upon cleavage of the GPI linkage (Siebold et al., 2017), we attempted to see, by high resolution *in situ* hybridization using RNAscope, which cochlear cells secreted the protein. RNAscope showed that RGMa mRNA was present in supporting cells that surround the nerve endings, suggesting that the protein was made in supporting cells and released to bind to its receptor on neurons (Fig. 1B). To add to our previous observation that neogenin1, the receptor for RGMa, was expressed in spiral ganglion neurons of the cochlea, we performed staining for neogenin1 and confirmed this localization in the nerve endings on the inner hair cells (Fig. 1C). Thus, both proteins are present at the synapse and their expression pattern suggests that RGMa released from supporting cells could bind to neogenin1 to repulse spiral ganglion neural growth cones at the inner hair cells.

**Figure 1.**
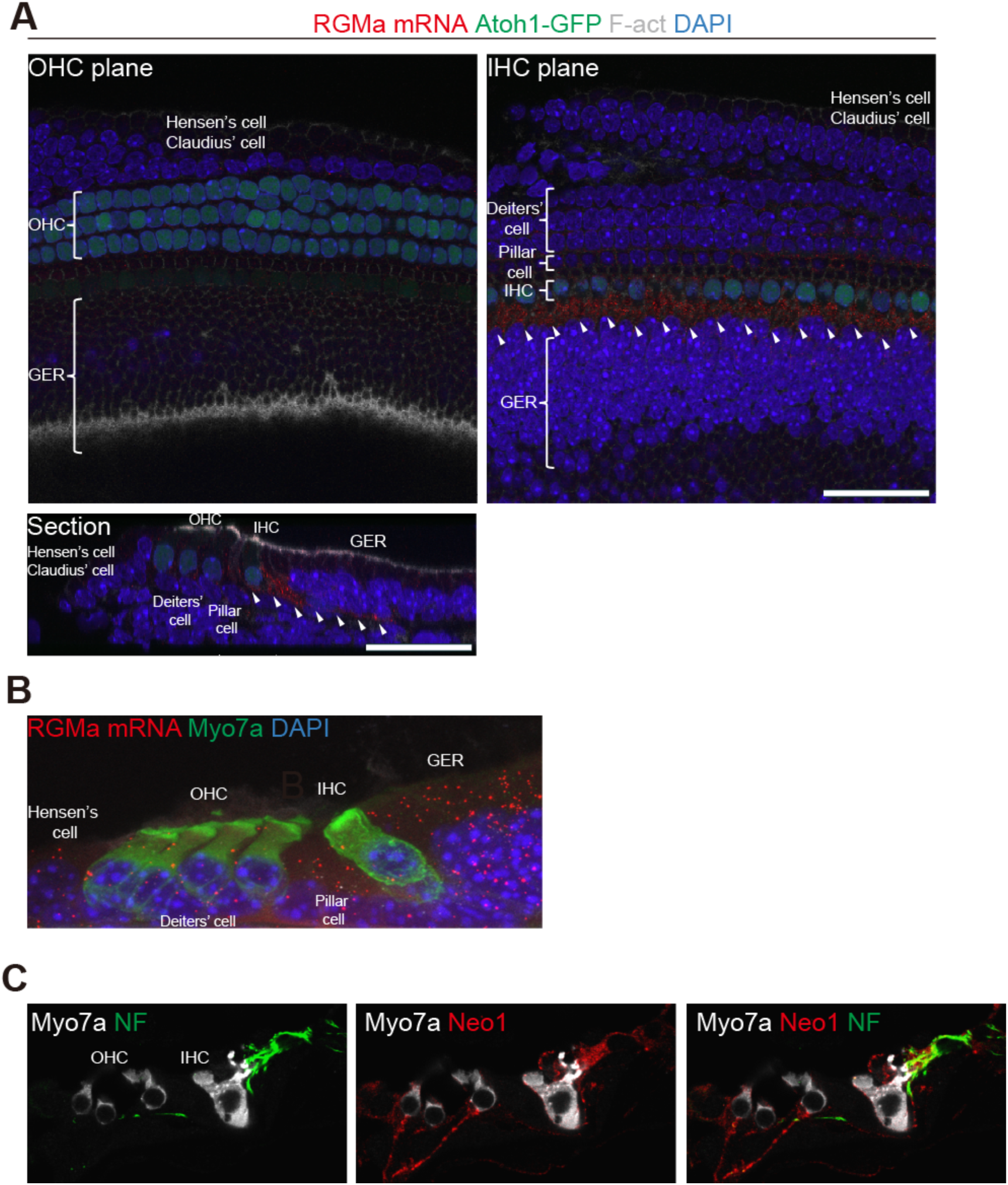
Localization of neogenin1 and RGMa in the mouse cochlea. (A) RGMa (white arrowheads) is detected by *in situ* hybridization (RGMa mRNA) in the supporting cells but not the hair cells of the P4 mouse organ of Corti. Hair cells are detected by phalloidin (f-actin) and Atoh1-GFP. (B) RGMa detection in a section of the organ of Corti shows mRNA in the pillar and Deiters’ cells and greater epithelial ridge (GER). (C) Neurofilament-positive afferent nerve fibers from spiral ganglion neurons of the adult cochlea reach the base of inner and outer hair cells (IHC and OHC). Neogenin1 is seen on the nerve terminals. Merged image shows co-labeling by neurofilament and neogenin1 antibodies. NF: neurofilament; Myo7a: myosin 7a; Neo1: neogenin1.

### Disruption of neonatal cochlear synapses by KA is reversed by treatment with anti-RGMa antibody

The postulated mechanism for disruption of synapses by noise involves excessive release of glutamate into the synaptic cleft with attendant damage to the postsynaptic neurons (Kujawa and Liberman, 2015). We used an *in vitro* model of excitatory cochlear synaptopathy to further probe the mechanism of the regenerative effect of the drug on cochlear synapses. In this model, cochlear explants, consisting of hair cells and attached spiral ganglion neurons, are exposed to KA, a selective ionotropic glutamate receptor agonist. The terminal processes of spiral ganglion neurons on hair cells are lost after KA exposure due to excitotoxic lesioning of the synapses in the cochlea (Pujol et al., 1985; Pujol and Puel, 1999). The time course of the model as described in the rat (Wang and Green, 2011) was reproduced in mouse ears; synapses were detected by the occurrence of CtBP2-expressing synaptic ribbons and PSD95-positive post-synaptic densities. After 2 h of treatment with KA, compared to control explants without KA treatment (Fig. 2A), damage was extensive at 5 h and reinnervation of hair cells by the peripheral processes of the neurons was apparent at 24 h (Fig. 2B). Growth was complete before 72 h, as the processes did not produce further reinnervation at this time point, and the peripheral processes were reduced in number in both the KA-treated and control samples, suggesting some deleterious effect of the prolonged culture.

**Figure 2.**
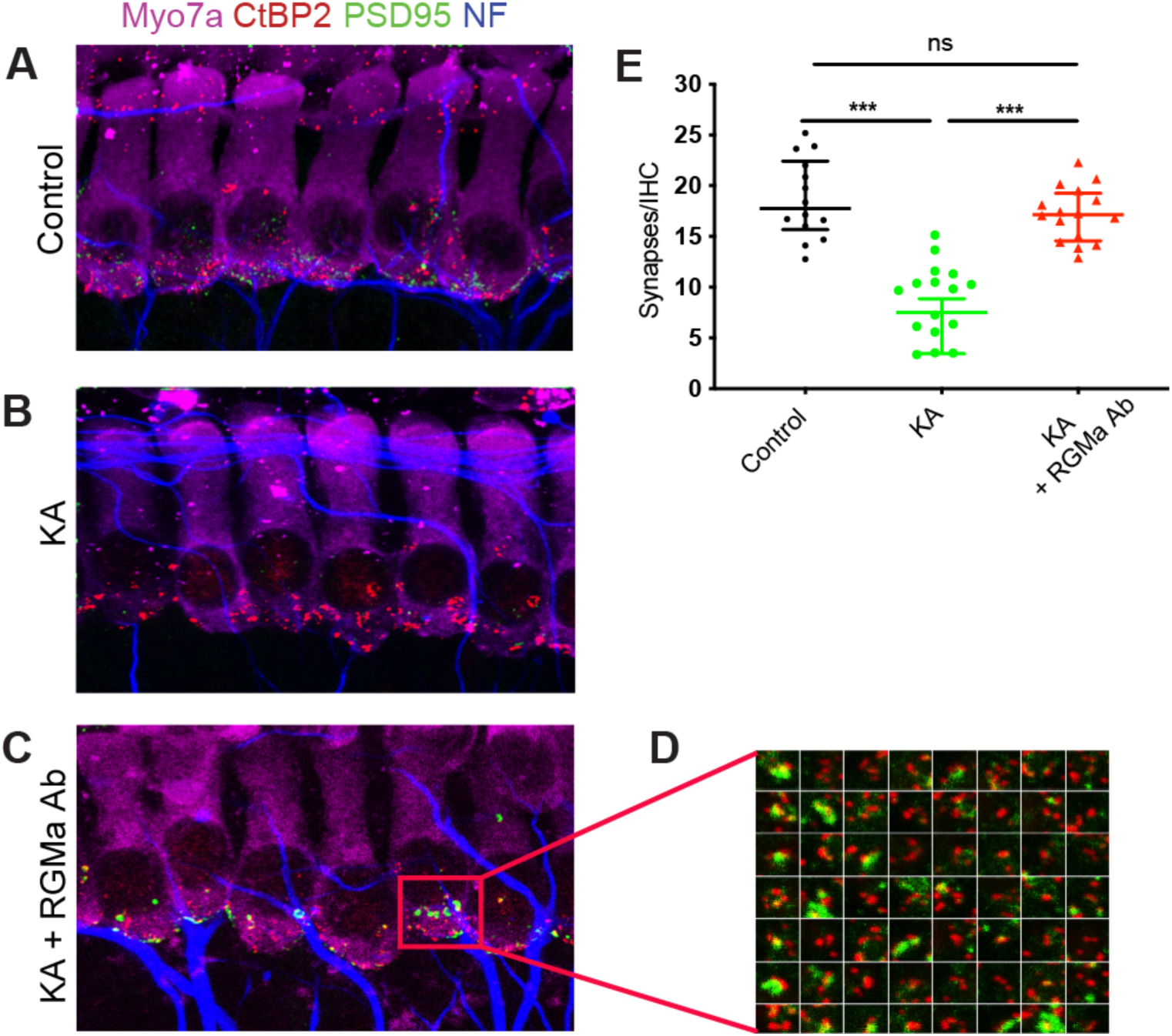
Disruption of neonatal cochlear synapses by kainic acid is reversed by treatment with anti-RGMa antibody. (A-C) Effects of kainic acid (KA) and RGMa antibody (RGMa Ab) on synapses between inner hair cells and auditory nerve fibers. Synaptic puncta are identified by juxtaposed label for PSD95 and CtBP2. Myosin 7a (Myo7a) is a marker for hair cells. (D) High-power thumbnails of a selection of paired-synaptic puncta. (E) The number of synapses significantly increased after kainic acid and anti-RGMa compared to kainic acid treatment alone. n=6, no treatment (Control); n=5, kainic acid only (KA); n=10 anti-RGMa (KA + RGMa Ab); ***: p < 0.001.

Treatment with anti-RGMa antibody in culture medium for 24 h after explant culture for 2 h with KA led to regeneration of synapses (Fig. 2C). Compared to IgG, the number of synapses was significantly increased in the anti-RGMa antibody treated cultures (Fig. 2D and E). Thus, in the glutamate toxicity model, anti-RGMa restored synaptic number and the restoration was specific to the anti-RGMa antibody.

### ABR thresholds recover 3 weeks after noise trauma

We next set out to test the effect of anti-RGMa antibody on synaptic loss in adult mice after noise exposure. Prior work defined noise parameters that caused synaptopathic hearing loss in the basal half of the murine cochlea (Jensen et al., 2015; Kujawa and Liberman, 2009, 2015; Suzuki et al., 2016). Loss of cochlear synapses is assessed by both counting of the afferent synapses at the base of inner hair cells and by analyzing the electrical signal at the level of the brainstem, where the first-order neurons of the auditory system, the spiral ganglion neurons, transmit the signal elicited by sound in the mechanotransducer cells, the hair cells. The electrical signal from the spiral ganglion neurons, which is transmitted to the brainstem and then up the central auditory pathway to the cortex, can be detected as a summed activity in the first wave of the auditory brainstem response (ABR). Exposure to a 2 h, 98 dB SPL octave band noise (8-16 kHz) results in a robust, but reversible threshold shift, as evidenced by DPOAE and ABR measures. A reduction in wave-I amplitude, without permanent alteration in the sound level required to elicit the response, and without loss of hair cells, as detected by the distortion product otoacoustic emissions (DPOAE), is the electrophysiological hallmark of cochlear synaptopathy. The noise causes no acute loss of inner or outer hair cells but initiates an immediate and permanent loss of synapses between auditory nerve terminals and cochlear inner hair cell ribbons in moderate to high frequency locations along the basilar membrane (Fernandez et al., 2015; Kujawa and Liberman, 2009, 2015; Sergeyenko et al., 2013).

We elicited cochlear synaptopathy by noise exposure to evaluate whether administration of anti-RGMa antibody after noise exposure ameliorated some of the irreversible damage caused by this noise exposure. Mice were treated, by round-window surgery, with anti-RGMa antibody or IgG, one week after noise exposure. Despite significant elevation at 24 h, consistent with previous reports (Liberman and Kujawa, 2017; Suzuki et al., 2016), ABR and DPOAE (Fig. 3A,B) thresholds were unaffected 3 weeks after exposure to 8-16 kHz octave-band noise at 98 dB SPL for 2 h (Fig. 3C). ABR thresholds are not permanently affected because the synapses remaining after synaptopathic noise respond normally to increasing sound level. DPOAE thresholds are not permanently affected because outer hair cells are not damaged at this level of noise exposure.

**Figure 3.**
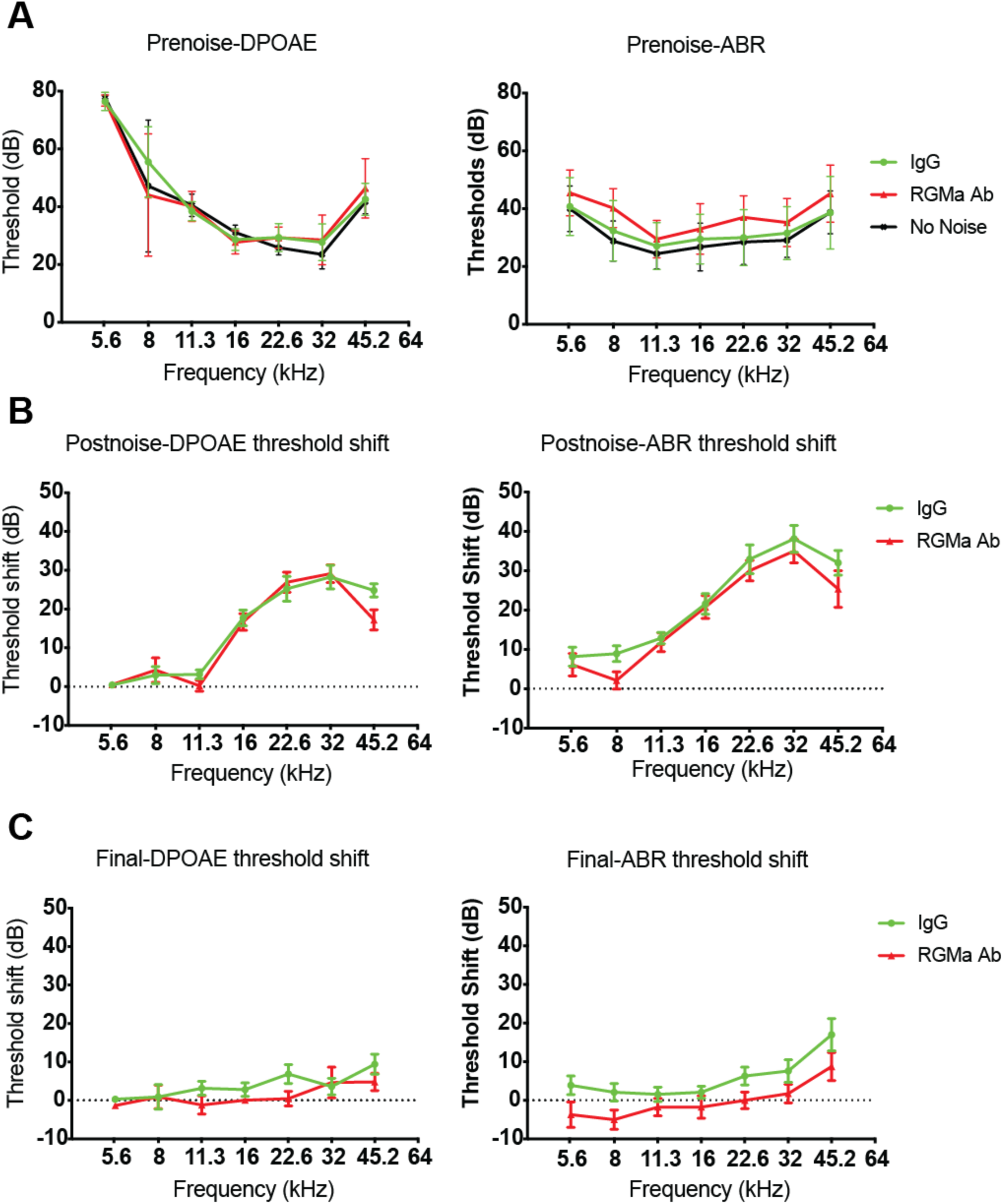
Threshold recovery 3 weeks after noise trauma. (A) DPOAE and ABR thresholds in unexposed (No noise, n=4), noise-exposed and anti-RGMa treated (RGMa Ab, n=15), and noise-exposed and IgG-treated (IgG, n=13) before noise trauma. (B, C) Post-noise and final DPOAE and ABR threshold shifts are defined relative to the mean thresholds for the same animals, measured respectively 24 h after the noise exposure, and before sacrifice i.e. 3 weeks after noise exposure. Temporary threshold shifts (TTS) are apparent at higher frequencies, post-noise, in all groups, without correlation to treatment. Error bars indicate SEM.

Treated animals with similar thresholds prior to the exposure (Fig. 3A) displayed as much as a 35-dB threshold elevation, relative to previously collected normative data, at test frequencies > 12 kHz through 44 kHz by both DPOAE and ABR (Fig. 3B), regardless of their assignment to the IgG or anti-RGMa treatment group. Repeat DPOAE and ABR (Fig. 3C) 3 weeks after exposure confirmed complete recovery of auditory thresholds for all groups.

### Recovery of wave-I amplitude and afferent synapse in anti-RGMa treated group

In comparison to threshold changes following noise, an analysis of suprathreshold ABR wave-I amplitudes showed no significant differences between the IgG-treated and anti-RGMa antibody-treated mice in the 11 kHz region (Fig. 4A). However, in the area of maximal temporary threshold shift, 32 kHz, a significant increase in amplitudes relative to IgG treated mice was achieved (p<0.01) for mice treated with anti-RGMa antibody (Fig. 4B). On average, amplitudes were 1.3 times larger for the anti-RGMa-treated vs the IgG-treated mice after the injection.

**Figure 4.**
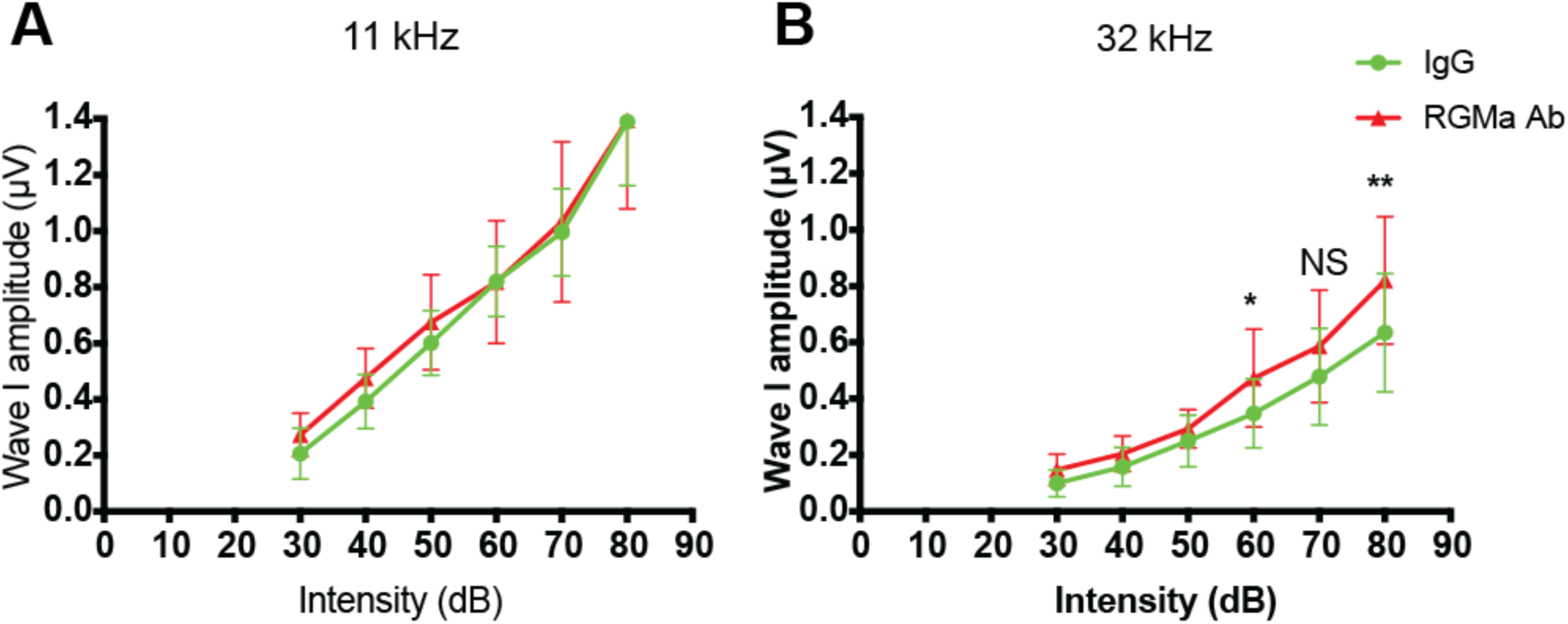
Recovery of wave 1 amplitude in anti-RGMa treated group. (A) Mean amplitude of wave-I in ABR 3 weeks after noise trauma i.e. 2 weeks after treatment by anti-RGMa or IgG, for each intensity of acoustic stimulation (dB SPL). There is no difference between the anti-RGMa (RGMa Ab) and control (IgG) groups at 11 kHz, where there is no synaptopathy. (B) Wave-I amplitude at 32 kHz increased 3 weeks after noise trauma in the anti-RGMa (RGMa Ab) as compared to the IgG group, indicating an effect of the antibody treatment. Statistical significance: *: p<0.05; **: p<0.01. Error bars indicate SEM.

As cochlear synapses are highly vulnerable to acoustic insult (Liberman et al., 2015), an assessment of the number of ribbon synapses per inner hair cell was conducted to determine if RGMa reversed synaptopathy. Relative to IgG-treated controls, anti-RGMa treated mice had comparable ribbon synapses/inner hair cell in the region with minimal noise-induced elevation (11 kHz, Fig. 5A). Yet significantly more synapses were seen in the 32 kHz region for mice receiving anti-RGMa (Fig. 5A, p<0.0001). The number of ribbon synapses/inner hair cell was approximately 1.5 times greater than the IgG group at 32 kHz. Immunohistochemistry for synapses showed an increase when treated with anti-RGMa at 32 kHz but no change at 11 kHz, consistent with the observation that the wave-I amplitude was increased at 32, but not at 11kHz (Fig. 5B). The increase in synapses could be seen in the 32 kHz region of the cochlea by comparison of the CtBP2/GluA2 pre- and post-synaptic staining pairs in the anti-RGMa treated as compared to the IgG-treated cochlea (Fig. 5C). Thus, 3 weeks after exposure, both ABR wave-I amplitudes and synaptic counts measured at 32 kHz were increased in the RGMa antibody-treated cochlea.

**Figure 5.**
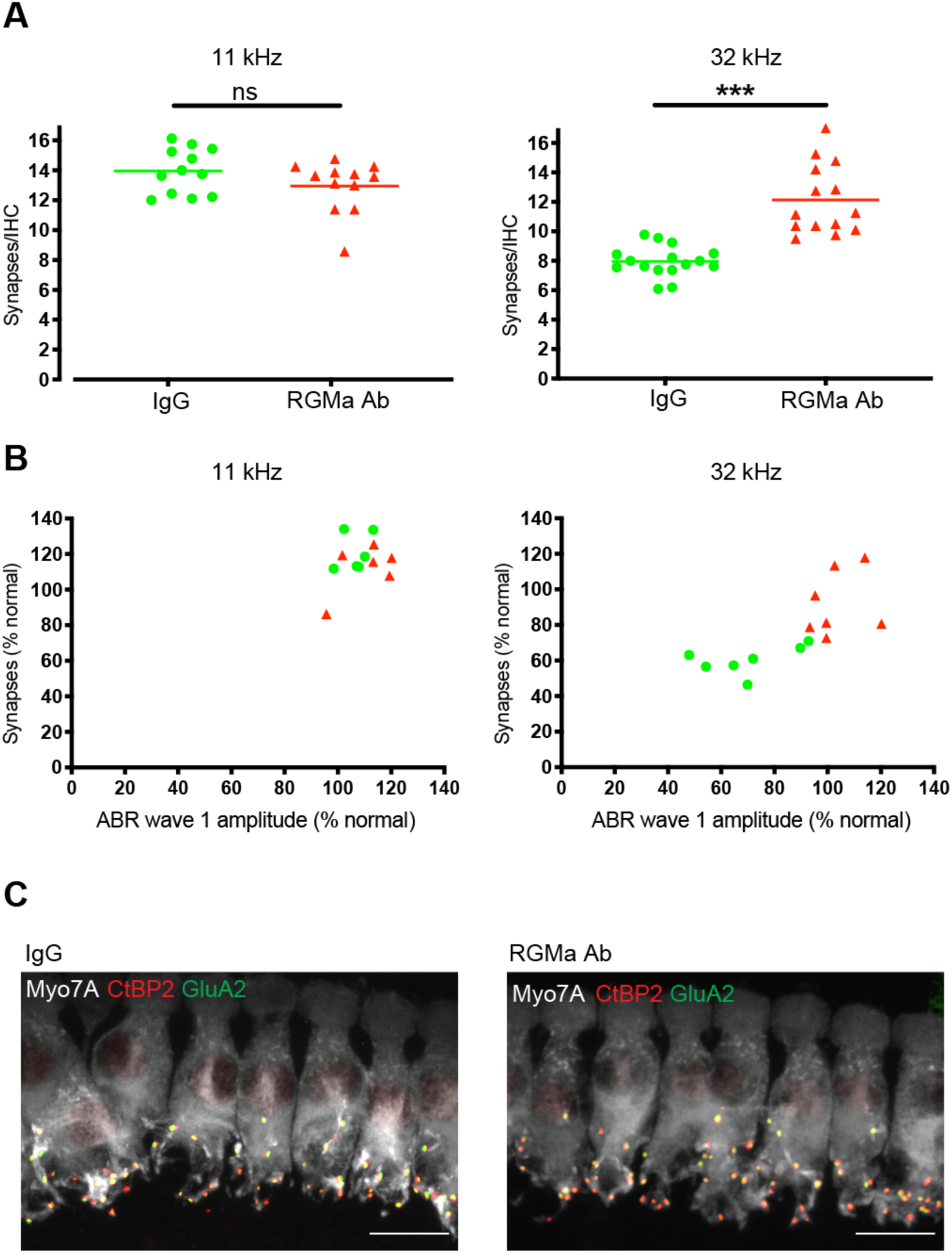
Number of ribbon synapses on inner hair cells are increased in anti-RGMa treated group. (A) Analysis of mean number of ribbon synapses in inner hair cells at 32 kHz shows a significant increase in the anti-RGMa treated group (11.7 ±3.27; n=10) compared to the IgG treated group (8.3 ±1.52; n=12). Red and green lines represent the mean number of synapses in each group. Statistical significance ***: p<0.001. (B) Correlation of wave-I amplitudes and synaptic counts at 11 and 32 kHz. (C) 3D confocal images of inner hair cells reveal ribbon synapses (CtBP2-positive) located at the base of the inner hair cells (myosin7a-positive). Ribbon synapses were considered orphan ribbons when not paired with post-synaptic receptor (GluA2-positive). CtBP2: C-terminal binding protein 2; GluA2: glutamate receptor 2; Myo7A: myosin 7a.

## DISCUSSION

Once established, synaptic loss at the primary afferent synapse is permanent, and there are no available treatments for hearing loss due to cochlear synaptopathy. The irreversibility of this clinical problem can be ascribed to the lack of regeneration of cochlear cells including the auditory neurons. Studies of synaptopathy have come to the conclusion that the disruption of this synapse by exposure to levels of noise that do not kill hair cells, nonetheless results in synaptic loss which does not recover (Kujawa and Liberman, 2006, 2009; Suzuki et al., 2016), even though the initial elevation of ABR threshold does partly recover. The lack of recovery of adult cochlear synapses after synaptopathic noise exposure can be documented by both physiological measures, the decreased amplitude of wave-I of the ABR, as well as by synaptic counts, which remain at the low level seen immediately after loss (Liberman and Kujawa, 2017).

Neurotrophins are required for development of the cochlear innervation and have been shown to protect neurons from damage due to toxins or other insults (Farinas et al., 1994; Farinas et al., 2001; Miller et al., 1997). Supporting cells secrete brain-derived neurotrophic factor (BDNF) and neurotrophin-3 (NT3) (Stankovic et al., 2004) during development and ablation of erbB in supporting cells results in aberrant innervation due to the lack of these secreted proteins. NT3 can regenerate cochlear synapses after acoustic overexposure when delivered locally in mice (Suzuki et al., 2016). We previously showed that a repulsive axonal guidance molecule was present in the cochlea (Brugeaud et al., 2014). Consistent with our previous description of the effect of anti-RGMa on neurite growth and synaptogenesis (Brugeaud et al., 2014), recovery from synaptic damage indicated an effect of RGMa antibody on the newborn cochlear synapse in a glutamate toxicity model, which is thought to be the pathophysiological mechanism for synaptic loss after exposure to noise (Kujawa and Liberman, 2015). The effect on the newborn was also seen in the adult, where treatment of noise-exposed animals with anti-RGMa one week after noise exposure also restored the cochlear afferent synapse and resulted in recovery of ABR wave-I amplitudes, indicating effective reversal of synaptopathy. This reversal is remarkable given the apparent inability of NT3 to effect the same recovery when administered after the insult (Hashimoto et al., 2019), and suggests that RGMa blocks access of growing fibers to the hair cells. Whether the reversal could be achieved at longer times remains unknown and will require us to characterize the latest effective time point of anti-RGMa administration after synaptopathic noise exposure. There is potentially a long therapeutic window for the treatment of synaptopathy because spiral ganglion neurons survive for months (in animal models) and years (in humans) after the causative insult. However, determining the most appropriate dose and mode of administration of the antibody would require further investigation.

RGMa is an axonal guidance molecule and like netrins and ephrins has primarily been studied in the developing nervous system; we and others have recently showed that RGMa and neogenin1 are expressed in postnatal tissue (Brugeaud et al., 2014). Neogenin1 and RGMa are upregulated after damage (Hata et al., 2006; Schwab et al., 2005). Upregulation of RGMa in spinal cord injury results in a block to regrowth of motor neurons, but treatment with anti-RGMa antibody could overcome this inhibition (Hata et al., 2006). Synaptic loss is thought to play a role in declining cognitive function in Alzheimer’s disease (Hong et al., 2016; Werneburg et al., 2020), and anti-RGMa had an effect on neural growth and connectivity in multiple sclerosis and spinal cord injury (Demicheva et al., 2015; Hata et al., 2006). Intracellular signaling elicited by neurotrophins is mediated in part by protein prenylation (Li et al., 2013), suggesting a link between neurotrophins and RGMa which has been shown to act through a prenylation pathway (Siebold et al., 2017).

The reversal of synaptic loss in the sensory neurons by anti-RGMa may have implications for other neurodegenerative disorders involving synaptic loss, as well as for an understanding of mechanisms of synaptic modulation important for neural plasticity. The effect on cochlear synaptic loss indicate that anti-RGMa could be used for the treatment of sensorineural hearing loss with cochlear neuropathy and could thus have translational implications for hearing loss. Hidden hearing loss results in compromised ability to understand words in a noisy background and has been implicated in tinnitus and hyperacusis. Further development of a treatment is therefore warranted and will require studies to determine optimal dosing after synaptopathic noise exposure either as a discrete event or a lifetime of smaller insults.

## Acknowledgements

Supported by NIH grant DC007174 (AE), Fondation pour l’audition (JN, MA), Fondation Philipps (JN), Fondation Monahan (JN), Fulbright fundation (JN), Fondation des Gueules Cassées (TB, LT), Audika (JN), Grand Audition (JN, LT), Association Française d’Oto-Neurologie (JN).

